# Analyzing Enzyme Kinetic Data Using the Powerful Statistical Capabilities of R

**DOI:** 10.1101/316588

**Authors:** Carly Huitema, Geoff Horsman

## Abstract

We describe a powerful tool for enzymologists to use for typical non-linear fitting of common equations in enzyme kinetics using the statistical program R. Enzyme kinetics is a powerful tool for understanding enzyme catalysis, regulation, and inhibition but tools to perform the analysis have limitations. Software to perform the necessary nonlinear analysis may be proprietary, expensive or difficult to use, especially for a beginner. The statistical program R is ideally suited to analyzing enzyme kinetic data; it is free in two respects: there is no cost and there is freedom to distribute and modify. It is also robust, powerful and widely used in other fields of biology. In this paper we introduce the program R to enzymologists who want to analyze their data but are unfamiliar with R or similar command line statistical analysis programs. Data are inputted and examples of different non-linear models are fitted. Results are extracted and plots are generated to assist judging the goodness of fit. The instructions will allow users to create their own modifications to adapt the protocol to their own experiments. Because of the use of scripts, a method can be modified and used to analyze different datasets in less than one hour.

## Introduction

Enzyme kinetics is a powerful tool for understanding enzyme catalysis, regulation, and inhibition. For example, details about transition state structures garnered from kinetic isotope effects on steady-state rate constants continues to inform the design of powerful transition state analog inhibitors of therapeutic value (Schramm, 2015, and Namanja-Magliano, Stratton, and Schramm, 2016). Moreover, the pioneering work of Cleland and many others over the last four decades has democratized enzyme kinetics so that it is no longer necessarily confined to specialists (Cook and Cleland, 2007). However, this popularization beyond the specialist community has driven demand for non-linear regression analysis software that is accessible to the non-specialist but also powerful enough to satisfy more advanced analyses. Although several commercial mathematical software packages exist, their costs can be prohibitive for labs in developing nations or other cost-conscious researchers. Standard software packages such as Excel can be adapted for these purposes (Kemmer and Keller, 2010), but the proprietary nature of the software makes customization difficult and presents a barrier to open science. Furthermore, spreadsheets are notorious for easy calculation mistakes and they are difficult to error check, test and validate because code is hidden away in tens or even hundreds of different cells. Proprietary software can also be a problem when expertise with one particular software tool becomes obsolete after moving to another organization that uses a different software package. Web-based tools offer a poor and partial solution, as they can be difficult to verify and transient in nature; a webpage used today may not be around tomorrow and a user has no ability to judge if the implementation is without error.

The statistical program R is an incredibly robust and powerful program that is widely used in life sciences for applications ranging from basic model fitting to large-scale analysis of gene expression data. It is open source, free to use, and available for multiple operating systems (Windows, Mac and Linux). From a training perspective, effort spent learning how to work in the R environment represents an easily transferable skill. However, with the robustness and flexibility of R comes a barrier to entry, as researchers may feel intimidated by its command line nature and lack of a user-friendly interface. Herein we provide a step-by-step guide for analyzing enzyme kinetic data in R, with the goal of increasing the accessibility to enzymologists of this powerful and free program. For the reader’s convenience we have included scripts as downloadable files in the supplemental online material.

## Getting started with R

The R program (a GNU project) can be acquired from the project webpage https://www.r-project.org/ by following the download and installation instructions for your specific operating system. Work in R is done using a command line and this program is sufficient for all analyses presented here. R Studio is a free and open-source integrated development environment for R. The base program R remains the same and work is still done using the command line. However, additional features (such as script integration and variable display) are added for easier development. There are many excellent tutorials and courses available online and in textbooks for an interested reader to get started using R (an excellent resource is the manuals available on the R project page: https://cran.r-project.org/). Here only the most basic introduction is given to provide the reader with the necessary starting point to perform the analyses described in this paper.

## Methods of inputting data into R

When R is started the user is presented with a prompt “>” and can enter commands. R can be used as a calculator, for example:

~~~
2^4 * 10 + log10(100)
~~~

gives a result of:

162

We can also use R to print out lists of numbers, for example:

~~~
−5:5 # anything after a pound/hash symbol is a comment and is ignored by R
~~~

gives a result of:

~~~
−5 −4 −3 −2 −1 0 1 2 3 4 5
~~~

An easy trick for lists is to multiply or divide the result:

~~~
−5:5/10
~~~

gives a result of:

~~~
−0.5 −0.4 −0.3 −0.2 −0.1 0.0 0.1 0.2 0.3 0.4 0.5
~~~

In R you can also work with variables that have been assigned values. The “=” symbol can be used to assign the value to a variable. However, the symbol “<-” is the more general assignment symbol than the “=” symbol and to avoid using both assignment symbols in this paper only “<-” will be used here.

~~~
a <- 2
~~~

*a* (not *A* as R is case sensitive) has been assigned the value of 2. Now the variable “*a*” can be used in calculations, for example:

~~~
a * 10
~~~

gives the result:

20

In evaluating enzyme kinetics, two of the most useful formats of storing data are in vectors and data frames. For example, the different concentrations of substrate used in an experiment can be stored in the vector *conc.uM*, using the function c() where c is short form for concatenate or paste together. The same can be done with the rate results.

~~~
conc.uM <- c(0.5,1,2.5,3.5,5,7.5,10,15,25,50,70,75,100)
rate <- c(0.6,1.1,2.1,2.3,3.7,3.,4.3,4.8,5.3,6.0,5.1,5.7,5.8)
~~~

The up arrow cycles through previously typed commands and can be useful to change something if you have made a mistake.

In enzyme kinetics the best way to organize experimental data in R is with a very important and useful data structure called a data frame. Data frames are used for storing tables of data consisting of columns of values. In a data frame the columns are named (for example conc.uM and rate) and can be referred to by name. To put our two vectors into a data frame called *exp1.df* we use the function data.frame().

~~~
exp1.df <- data.frame(conc.uM, rate)
~~~

and to look at the contents of our data frame simply type the variable name:

~~~
exp1.df
~~~

To access a column by name we use the $ symbol:

~~~
exp1.df$conc.uM
~~~

If you have a very large data frame you may only want to look at a portion of it to check the structure of the data. The functions head() or tail() will show you the first or last 6 entries in the data frame as well as the labels of the columns.

~~~
head(exp1.df)
~~~

To count the number of rows in the data frame and confirm all your data is present use the function nrow().

~~~
nrow(exp1.df)
~~~

We can perform different operations on the data in the data frame including adding new data. For example, before we perform further analysis we may want to work with the concentration in nM instead of μM. We can add another column to the data frame where we multiply *exp1.df$conc.uM* by 1000 to give us concentrations in nM with the command:

~~~
exp1.df$conc.nM <- exp1.df$conc.uM * 1000
~~~

If you look at the contents of the data frame *exp1.df* now you will see the addition of a third column named conc.nM with the concentrations now expressed in nM.

We have been creating objects in R and we can see all the objects that have been created in the current R session by using the function ls().

~~~
ls()
~~~

To remove all these objects we have created we use the command rm(list=ls()). Briefly, the command rm(list=“”) will remove all objects passed (using the “=” sign) to the argument “list”. If we pass all objects in the R environment with the function ls(), we will remove all objects.

~~~
rm(list=ls())
~~~

You can also remove specific objects using this command, for example with rm(list=c(“conc.uM”, “rate”)).

Data can be added manually to R using the functions c() and dataframe(), however, it can be easier to input the data from a .csv file which can be the output of many programs including Excel.

The read.csv() function reads data from a .csv file and puts it directly in a data frame. It assumes that the first row specifies the names of your columns but if this is not the case you can add the argument “header=FALSE” to the function.

~~~
exp1.df <- read.csv(“filename.csv”, header=TRUE)
~~~

To know which folder/directory that R is looking in for your data file (your current working directory) use the function getwd().

To avoid the challenge of different computers with different working directories, in all examples in this paper the data will be inputted directly rather than read from a file. In this manner, all the scripts are complete and no extra data files are needed.

Finally, rather than typing the commands into R directly each time analysis is performed, the commands can be put in a simple text file (the script file) using an editor such as notepad in the Windows environment, and then copied and pasted directly into R. In this way, the commands used to generate the data can be saved and then put with the data as a record of analysis, and copied and modified with each new experiment. After gaining familiarity with R, users are encouraged to learn the feature rich R Studio program, which better integrates scripts into the R environment.

### Plotting data

Data plotting in R is robust, with many examples available on the internet demonstrating all that R is capable of. However, here we will only describe the most basic plotting techniques that will be useful for fitting enzyme kinetics. The reader may decide if they prefer to export the data and prepare figures in their chosen program or to learn more about plotting in R from other resources.

Before any analysis of an enzyme kinetic experiment, the experimental results should be examined on a plot using the function plot(x,y) where *x* is the vector of values for the x axis and *y* the vector of values for the y axis. Using this dataset:

~~~
conc.uM <- c(0.5,1,2.5,3.5,5,7.5,10,15,25,50,70,75,100)
rate <- c(0.6,1.1,2.1,2.3,3.7,3.,4.3,4.8,5.3,6.0,5.1,5.7,5.8)
~~~

We can plot rate vs concentration with the command:

~~~
plot(conc.uM, rate)
~~~

In this example, the x and y axis will have labels with the name of the variables. To have our own custom title and axis labels we add a few arguments to the plot() function with the command:

~~~
plot(conc.uM, rate, main=“Plot Title”, xlab=“conc (uM)”, ylab=“rate (uM/min)”)
~~~

It is also useful to add lines to our plot, perhaps the line of best fit after we have fit our model to the data. We can add lines to our plot with the function lines(x, y, lty, col) where lty is line type and col is the color of the line.

~~~
lines(conc.uM, rate, lty=“dotted”, col=“red”)
~~~

For this example, we are simply joining the points of data, which is not very useful. However, once we have performed a fit we can generate a set of theoretical x, y data points using the solved parameters and use them to draw a smooth line on our plot showing our fit.

### Fitting the Michaelis-Menten equation

The Michaelis-Menten equation is the fundamental equation of enzyme kinetics (Bowden, 2004) and has the typical format of:

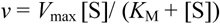

*V_max_* is the maximum theoretical reaction rate where additional substrate does not noticeably increase the substrate turnover (*V*_max_ is theoretical because it is at the asymptote that the fitted curve will never reach). *K_M_* is the Michaelis constant and often oversimplified in being described as the affinity of the enzyme for the substrate. The Michaelis constant is always the concentration of substrate ([S]) where the rate *v* is half that of *V_max_*.

The plot of results from an experiment to determine *K_M_* and *V_max_* has substrate concentration [S] on the x-axis and reaction rate *v* on the y-axis. Prior to the era of personal computing it was acceptable to transform this data into a linear format (typically the Lineweaver-Burk plot) in order to generate a fit for the data and determine *K_M_* and *V_max_*. However, today it is recommended to fit the Michaelis-Menten equation directly to the data using nonlinear least-squares fitting.

An excellent introduction to nonlinear least-squares fitting and how to perform this fitting in Excel is described by Kemmer and Keller (2010). Unlike linear regression there is no analytical solution to the fitting, instead a fit is determined through trial and error. The fitting algorithm begins with estimated starting values and attempts to lower the sum of square of errors (the difference between expected and calculated values, squared) by altering the parameters until a minimum is reached (or the algorithm cannot find a solution). In this process there is no guarantee that a global minimum has been obtained. As only a local minimum is found, a good estimate of starting parameters is important.

Here are the commands for performing a fit followed by detailed instruction and description.

~~~
# Fit Michaelis-Menten equation
# input the data and plot
conc <- c(0.5,1,2.5,3.5,5,7.5,10,15,25,50,70,75,100)
rate <- c(0.6,1.1,2.1,2.3,3.7,3.,4.3,4.8.5.3.6.0.5.1.5.7.5.8)
test.df <- data.frame(conc,rate)
plot(test.df$conc,test.df$rate)
~~~

~~~
# perform the fitting
mm.nls <- nls(rate ~ (Vmax * conc / (Km + conc)), data=test.df,
start=list(Km=5, Vmax=6))
summary(mm.nls)
~~~

~~~
# extract coefficients
Km <- unname(coef(mm.nls) [“Km”])
Vmax <- unname(coef(mm.nls)[“Vmax”])
~~~

~~~
# plot data and plot line of best fit
x <- c(0:100)
y <- (Vmax*x/(Km+x))
lines(x,y, lty=“dotted”, col=“blue”)
~~~

~~~
# confidence intervals of parameters confint(mm.nls)
~~~

~~~
# look at residuals and plot
mm.resid <- resid(mm.nls)
plot(test.df$conc, mm.resid)
~~~

~~~
# add weighting to fit
test.df$weight <- 1/test.df$conc^2
mm.weight.nls <- nls (rate ~ (Vmax *
conc / (Km + conc)), data=test. df,
start=list(Km=5,       Vmax=6),
weight=test.df$weight)
summary(mm.weight.nls)
~~~

To begin with fitting, we will start by entering data, putting it into a data frame and plotting the data to evaluate the potential for fitting using the commands:

~~~
conc <- c(0.5,1,2.5,3.5,5,7.5,10,15,25.50.70.75.100)
rate <- c(0.6,1.1,2.1,2.3,3.7,3.,4.3,4.8.5.3.6.0.5.1.5.7.5.8)
test.df <- data.frame(conc,rate) test.df
# have a look at your data
plot(test.df$conc,test.df$rate)
~~~

By examining the plot we can see that the data appears suitable for fitting a Michaelis-Menten curve and from the plot we can estimate initial starting values for the parameters *K_M_* and *V_max_*. In this example, good estimates of *V_max_* and *K_M_* would be 6 and 5 respectively.

In R, the function for fitting using nonlinear least- squares is nls() and there are several arguments that must be passed in the function for proper analysis. First, the command:

~~~
mm.nls <- nls(formula(rate ~ (Vmax * conc / (Km + conc))), data=test.df, start=list(Km=5, Vmax=6))
~~~

The formula to be fit is the first argument “formula(rate ~ (Vmax *conc/(Km + conc))”. *rate* and *conc* are variables that match the column names in the data frame *test.df* and *Vmax* and *K_M_* are parameters that we wish to solve for. We specify which data frame to be used in the fitting using the argument data=test.df. Finally, we must give the function starting values for the two parameters that we wish to solve with the argument start=list(Km=5, Vmax=6). The names of these variables and parameters must match either column labels in the data frame or listed in the starting values or R will serve the user an error. Note that the arguments are separated by commas. The result of the fitting is saved in the object mm.nls. It contains information from the fit including estimates of the parameters, test statistics, residuals etc.

We can add an additional argument trace=TRUE or FALSE to the function nls(), this will show us each iteration during the fitting and the different values of *Vmax* and *K_M_* that are evaluated and can be useful for troubleshooting if the fit is not successful. If the fitting fails one of the first things to test is better starting values. A fitting may also fail if the data being fit does not represent the model.

After the fitting is complete we can view a summary of the model including values determined for the parameters with the function summary ().

~~~
summary(mm.nls)
~~~

To see the fit on the plot with the data we first generate a vector of *x* values (concentration) covering the range we wish to plot. We use the function c() to create our vector with values from 0 to 100.

~~~
x <- c(0:100)
~~~

Then for each value we calculate the expected rate values for each of these concentrations using the values of the parameters revealed using the summary() function (*Vmax*=6.069 and *Km*=4.701).

~~~
y <- (6.069*x/(4.701+x))
~~~

Finally, we draw a line on the plot with this generated data giving us our line of best fit. The function lines() adds to an existing plot, so you must not close a plot window when you intend to add lines to your figure.

~~~
lines(x, y, lty=“dotted”, col=“red”)
~~~

Rather than copying the values of the parameters manually we can extract the coefficients from the model using the function coef().

~~~
coef(mm.nls)
~~~

We can access each coefficient by name (used within square brackets and quotes) and then assign the value to a new variable. Sometimes a named variable can cause problems when using other functions in R so we strip the name from the coefficient using the function unname().

~~~
Km <- unname(coef(mm.nls) [“Km”])
Vmax <- unname(coef(mm.nls)[“Vmax”])
~~~

Now we can use the values directly in the equation to calculate line of best fit:

~~~
y.2 <- (Vmax*x/(Km+x)) lines(x,y.2, lty=“dotted”, col=“blue”)
~~~

Once we have fit our model we can extract the confidence intervals for the parameters from the fit object *mm.nls* using the function confint().

~~~
confint(mm.nls)
~~~

Finally, after performing a fit it is important to plot the residuals from your fit to determine if the model selected is good and if the fit was successful. The values of the residuals can be accessed from the fit object *mm.nls* using the function resid().

~~~
mm.resid <- resid(mm.nls)
plot(test.df$conc, mm.resid)
~~~

A common technique when fitting experimental data is to weight the fit. To add weighting add another column to your data in the data frame with the values for the weighting. For example, to weight the low concentration values more in the fit we can use an equation of weight = 1/conc^^^2. The following command assigns these weighting values to a new column in the *test.df* data frame.

~~~
test.df$weight <- 1/test.df$conc^2
~~~

To perform nonlinear least-squares analysis with weighting we add a new weight argument to our nls() function.

~~~
mm.weight.nls <- nls(rate ~ (Vmax * conc / (Km + conc)), data=test.df,
start=list(Km=5, Vmax=6),
weight=test.df$weight)
~~~

Fitting Michaelis-Menten equation with substrate inhibition

The effect of substrate inhibition can be accounted for with a different model:

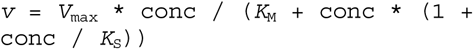

Where *K_S_* is the parameter that accounts for the inhibition of the substrate. However, fitting is very similar to the simple Michelis-Menten equation with just an additional parameter to the nls() fitting function.

~~~
# Fit Michaelis-Menten equation with
substrate inhibition
# input the data and plot
conc <- c(0.01,0.02,0.03,0.04,0.06,0.08,0.1,0.3,0.3,0.4,0.6,0.8,1,1,2,3,4,5,5,5,6)
rate <- c(3.1,5.2,5.6,5.9,7.2,7.6,8.4,9.2,10.4,10,11,10.9,10.3,10.4,10.1,9.6,9.4,9.6,8.6,8.7,8.5)
test.df <- data.frame(conc,rate)
plot(test.df$conc,test.df$rate)
# perform the fitting, look at plot to estimate start values
mminhib.nls <- nls(rate ~ (Vmax * conc / (Km + conc*(1+conc/Ks))),
data=test.df, start=list(Km=0.2,
Vmax=11, Ks=1))
summary (mminhib.nls)
# extract coefficients
Km <- unname(coef(mminhib.nls)[“Km”])
Vmax <-
unname (coef (mminhib.nls) [“Vmax”])
Ks <- unname(coef(mminhib.nls)[“Ks”])
# plot data and plot line of best fit
x <- c(0:60)/10
y <- (Vmax*x/(Km+x*(1+x/Ks)))
lines(x,y, lty=“dotted”, col=“blue”)
# confidence intervals of parameters confint (mminhib.nls)
# look at residuals and plot
mminhib.resid <- resid(mminhib.nls)
plot (test.df$conc, mminhib.resid)
~~~

### Determining IC_50_

The IC_50_ of an inhibitor is the concentration of an inhibitor where the enzyme activity is half of maximum and is done by fitting a four-parameter Scurve to the experimental data (in this example the 4-parameter logistic model (Rodbard and Frazier, 1975)) using the nls() function in R. First the script is presented followed by a description.

~~~
# Calculate IC50
# enter the data into a data frame and plot
conc.uM <- c(300,150,75,38,19,5,2,1,0.6)
percent.activity <- c(2,7,12,22,36,53,67,83,85)
ic50.df <- data.frame(conc.uM, percent.activity)
ic50.df$conc.nM <- ic5 0.df$conc.uM * 1000
ic50.df$logconc.nM <- log10(ic50.df$c onc.nM)
plot(ic50.df$logconc.nM,ic5 0.df$perce nt.activity)
# estimate initial values of the curve by examining the plot
# add a 4-parameter rodbard curve with these initial parameters to the plot
# to check they are reasonable
initial estimates
x <- c(2:12/2)
y <- 0+(80-0)/(1+(x/4)^^^10)
lines(x,y, col=“red”)
# fit the data using the nls()
function to the 4-parameter logistic model
rodbard.fit <- nls(formula(percent.activity ~ bot+ (top-
bot)/(1+(logconc.nM/logic50)^slope)),
algorithm=“port”, data=ic50.df,
start=list(bot=0, top=80, logic50=4,
slope=10), lower=c(bot=-Inf, top=-
Inf, logic50=0, slope=-Inf))
# generate a summary of the fit
summary(rodbard.fit)
# extract coefficients
top <- unname(coef(rodbard.fit) [“top”])
bot <- unname(coef(rodbard.fit) [“bot”])
logic50 <- unname(coef(rodbard.fit) [“ logic50”])
slope <- unname(coef(rodbard.fit)[“slope”])
# calculate line of best fit and add
to plot
y.fit <- bot+(top-bot)/(1+(x/logic50)
^slope)
lines(x,y.fit, col=“green”)
# log scale to linear scale to get
IC50
# remember the results are in nM
10^logic50
~~~

The 4-parameter logistic model is of the form *y* = *d* + *(a* - *d)* / (1 + (*x/c*)^^^*b*) where *x* is the log(conc), *y* is the inhibition (or percent activity) and *a*, *b*, *c* and *d* are parameters. The parameter *c* is the log(IC_50_) and is the parameter we are interested in. To estimate the initial starting values for using in the nls() function examine the plotted data. Estimates for *a* and *d* are the expected highest and lowest y-axis values of the fitted curve respectively and *c* the value of the x-axis at the midpoint of inflection of the curve. Parameter *b* controls the steepness of the curve and reasonable starting values of *b* are 10 if the curve curves down (if you are graphing percent activity) and -10 if the curve goes up (if you are graphing percent inhibition). Initial starting values are at best guesses and plotting expected curves with the data will help in refining the estimates.

It is important when fitting with the 4-parameter logistic model that the log of the concentrations (*x*) must all be positive (*x* values must all be greater than 1). The reason is mathematical, when *x* is negative and *b* is not a whole number (*x/c*)*^^^b* can result in a complex number and you will get an error when attempting to fit the model. In this example, the concentration of inhibitor used in the fit is converted to nM which results in no negative values of *x*.

### Fitting competitive inhibition

Competitive inhibition can be described by the model:

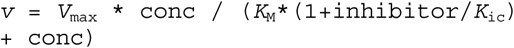

Fitting competitive inhibition is similar to the fitting of Michaelis-Menten data but with some more sophistication necessary to make a generally useful script; specifically, data subsets and loops. A script for fitting competitive inhibition is presented here and a description of the script follows.

~~~
# Fit competitive inhibition
# input the raw data into a data frame
conc <- c(3.9,1.9,0.7,0.5,0.4,0.3,3.9,1.9,0.7,0.5,0.4,0.3,3.9,1.9,0.7,0.5, 0.4,0.3,3.9,1.9,0.7,0.5,0.4,3.9,1.9,0.7,0.5,0.4)
rate <- c(0.19,0.19,0.18,0.14,0.13,0.08,0.18,0.17,0.15,0.13,0.11,0.11,0.2,0.15,0.11,0.1,0.09,0.07,0.16,0.14,0.1,0.08,0.07,0.16,0.13,0.08,0.08,0.07) inhibitor <- c(0,0,0,0,0,0,100,100,100,100,100,100,300,300,300,300,300,300,500,500,500,500,500,700,700,700,700,700)
kic.df <- data.frame(conc, rate, inhibitor)
# determine concentrations of
inhibitor used in experiment
inhibitor.conc <-
unique(kic.df$inhibitor)
# create colors for each inhibitor
concentration
inhib.color <-
rainbow (length (inhibitor.conc))
# generate a blank plot and then plot
the raw data
plot (kic.df$conc,kic.df$rate, pch=“”)
for (i in 1:length(inhibitor.conc)) {
   points(subset(kic.df$conc,
kic.df$inhibitor==inhibitor.conc[i]),
subset(kic.df$rate,
kic.df$inhibitor==inhibitor.conc[i]),
col=inhib.color[i], pch=1)
}
# perform the fitting
kic.nls <- nls(rate ~ (Vmax * conc /
  (Km* (1+inhibitor/Kic) + conc)),
data=kic.df, start=list(Km=0.5,
Vmax=.2, Kic=300))
# generate a summary of the fit
summary (kic.nls)
# confidence intervals of parameters
confint(kic.nls)
# extract coefficients
Km <- unname(coef(kic.nls)[“Km”])
Vmax <- unname(coef(kic.nls)[“Vmax”])
Kic <- unname(coef(kic.nls)[“Kic”])
# use the values directly in the
equation to calculate line of best
fit
fit.data <- expand.grid(x=(1:40)/10,
inhib=inhibitor.conc)
fit.data$y <- Vmax*fit.data$x/(Km*(1+
fit.data$inhib/Kic)+fit.data$x)
# plot lines of best fit
for (i in 1:length(inhibitor.conc)) {
   lines (subset (fit.data$x,
fit.data$inhib==inhibitor.conc[i]),
subset(fit.data$y,
fit.data$inhib= = inhibitor.conc[i]),
col=inhib.color[i])
}
~~~

The experimental data for competitive inhibition consists of measurements of reaction rates for different concentrations both substrate and inhibitor. The entire dataset is fit to the competitive model and so the data cannot be in a ‘wide’ form with a column for each inhibitor concentration. Instead, we will have a ‘long’ form dataframe with three columns: substrate concentration, inhibitor concentration and measured rate for each experimental measurement.

~~~
conc <- c(3.9,1.9,0.7,0.5,0.4,0.3,3.9,1.9,0.7,0.5,0.4,0.3,3.9,1.9,0.7,0.5, 0.4,0.3,3.9,1.9,0.7,0.5,0.4,3.9,1.9,0.7,0.5,0.4)
rate <- c(0.19,0.19,0.18,0.14,0.13,0.08,0.18,0.17,0.15,0.13,0.11,0.11,0.2,0.15,0.11,0.1,0.09,0.07,0.16,0.14,0.1,0.08,0.07,0.16,0.13,0.08,0.08,0.07)
inhibitor <- c(0,0,0,0,0,0,100,100,100,100,100,100,300,300,300,300,300,300,500,500,500,500,500,700,700,700,700,700)
kic.df <- data.frame(conc, rate,
inhibitor)
plot (kic.df$conc,kic.df$rate)
~~~

While this ‘long’ dataframe format is necessary for the fitting, a downside is obvious in a plot of this data; we do not visually distinguish which points correspond to different inhibitor concentrations. However, inhibitor concentrations are accounted for in the model fitting and making a more human readable plot will be covered at the end of this section.

As before, we can fit the competitive inhibition model using the nls() function, extract the coefficients and calculate confidence intervals of the parameters.

~~~
kic.nls <- nls(rate ~ (Vmax * conc / (Km* (1+inhibitor/Kic) + conc)), data=kic.df, start=list(Km=0.5, Vmax=.2, Kic=300))
summary (kic.nls)
Km <- unname(coef(kic.nls)[“Km”])
Vmax <- unname(coef(kic.nls) [“Vmax”])
Kic <- unname(coef(kic.nls)[“Kic”])
confint (kic.nls)
~~~

To plot our line of best fit in R we introduce the very useful function expand.grid() which creates a data frame using all combinations of supplied vectors (in this case *x* and *inhib*).

~~~
fit.data <- expand.grid(x=(1:40)/10, inhib=c(0,100,300,500,700))
~~~

Using the generated data we calculate rates (*y*) for the lines of best fit and graph the results on the raw data plot.

~~~
fit.data$y <- Vmax*fit.data$x/(Km*(1+
fit.data$inhib/Kic)+fit.data$x)
lines(fit.data$x,fit.data$y,
lty=“dotted”, col=“red”)
~~~

Looking at this example we can see the fit of the result but it is not so easy to interpret. We want to plot 5 different lines (one for each inhibitor concentration) but we instead plot one single line which awkwardly connects all the data at all inhibitor concentrations. For an initial quick look this may be sufficient.

For a more easily interpreted plot of our data we must take our large data frame and subset it and then plot lines for each inhibitor concentration separately. We can do this using the function subset() where the first argument in the function is the data to be subsetted (*fit.data*) and the second argument is the logical expression that is being evaluated (*inhib==0*). Watch for the double equal signs in the logical expression! The following commands pull out a subset of data where the concentration of inhibitor is zero, assigns it to a new data frame *fit.0* and plots the result.

~~~
fit.0 <- subset(fit.data, inhib==0)
lines(fit.0$x,fit.0$y, lty=“dotted”,
col=“blue”)
~~~

And then for another concentration of inhibitor:

~~~
fit.100 <- subset(fit.data,
inhib==100)
lines(fit.100$x,fit.100$y,
lty=“dotted”, col=“red”)
~~~

We can imagine that manually creating different data subsets for each concentration of inhibitor achieves the plotting result we want but this approach is time consuming and not a very general solution. Fortunately, with a bit more sophistication we can create a more general script using for loops.

Using our data frame *kic.df*, we want to identify all the unique concentrations of inhibitor are present using the function unique().

~~~
unique(kic.df$inhibitor)
~~~

We then store that value in the vector *inhibitor.conc*. Using the function length() we can see there are five values in *inhibitor.conc*.

~~~
inhibitor.conc <-
unique (kic.df$inhibitor)
length (inhibitor.conc)
~~~

Square brackets can be used to identify the n’th value in a vector. For example, the value of the third entry in the vector *inhibitor.conc* is 300.

~~~
inhibitor.conc[3]
~~~

We can substitute a variable within the square brackets (in this example *i*) to identify the same value.

~~~
i=3
inhibitor.conc[i]
~~~

Using our vector of inhibitor concentrations (*inhibitor.conc*) we can subset the data as we saw previously using the same subset() function.

~~~
i=3
subset(kic.df$conc,
kic.df$inhibitor==inhibitor.conc[i])
~~~

Then we can plot this subset of values and produce a plot with only one inhibitor concentration (300 in this example). The function plot is in the form plot(x,y, col) - look for the comma separating x and y vectors.

~~~
plot (subset (kic.df$conc,
kic.df$inhibitor==inhibitor.conc[i]),
subset (kic.df$rate,
kic.df$inhibitor==inhibitor.conc[i]),
col=“blue”)
~~~

There is one more tool that we need before we can make our plotting loop, the color. We want each different concentration in the plot to be shown with a different color. R can generate a vector of colors with the rainbow() function where the number in the brackets is the number of colors to be generated.

~~~
rainbow(3)
~~~

We can make a vector containing one colour for each inhibitor concentration in our data frame.

~~~
inhib.color <-
rainbow (length (inhibitor.conc))
inhib.color
~~~

And the rewritten plot() function with the color determined by rainbow() will be:

~~~
i=3
plot (subset(kic.df$conc,
kic.df$inhibitor==inhibitor.conc[i]),
subset(kic.df$rate,
kic.df$inhibitor==inhibitor.conc[i]),
col=inhib.color[i])
~~~

Finally, to plot our data we first create an empty plot of the dimensions that will fit all the data, then we will go through a loop and plot the rate vs concentration for each concentration of inhibitor. We could create the empty plot of the correct size using the function new.plot() but we will need to specify details such as axis size and labels manually. As a short-cut we can create the plot using all the data but with ‘invisible’ points (we use the argument pch=“”, that is, the point character ‘pch’ is nothing “”). This way a plot will be automatically created that will fit the entire dataset.

~~~
plot(kic.df$conc,kic.df$rate, pch=“”)
# our blank plot
~~~

Now all the components come together in a loop which loops 5 times (the length of the vector *inhibitor.conc*) and for each inhibitor concentration it plots the substrate concentration versus rate in a different color. To demonstrate the looping (and as a useful troubleshooting technique) the function print() will display values as the loop runs. In this example print(i) will return the value of *i* for each loop (1, 2, 3, 4, and 5). The function points() is similar to the function lines() only it plots points instead of lines.

~~~
for (i in 1:length(inhibitor.conc)) {
   print (i)
   points (subset (kic.df$conc,
kic.df$inhibitor==inhibitor.conc[i]),
subset(kic.df$rate,
kic.df$inhibitor==inhibitor.conc[i]),
col=inhib.color[i],
lty=0, pch=1)
}
~~~

The advantage of this loop method is that once a script is written, with the same few short lines of script different data sets can be displayed without requiring much extra effort.

To plot the lines of best fit (as calculated using the function nls() as described earlier in this example) we generate the data again using the function expand.grid() and this time instead of manually specifying the inhibitor concentrations we use the vector *inhibitor.conc*.

~~~
fit.data <- expand.grid(x=(1:40)/10,
inhib=inhibitor.conc)
fit.data$y <-
Vmax*fit.data$x/ (Km*(1+fit.data$inhib
/Kic) +fit.data$x)
~~~

Then we use another loop to draw lines of best fit for each concentration of inhibitor.

~~~
for (i in 1:length(inhibitor.conc)) {
   lines(subset(fit.data$x,
fit.data$inhib==inhibitor.conc[i]),
subset(fit.data$y,
fit.data$inhib==inhibitor.conc[i]),
col=inhib.color[i])
}
~~~

Script with Uncompetitive, Competitive and Mixed inhibition

This script uses all the techniques from this paper and fits data to uncompetitive inhibition, competitive inhibition and mixed inhibition models. It plots all the data and the fits as well as the residuals from the fitting. Finally, it plots the Lineweaver-Burk transformation of the data with fits for visual confirmation. No new techniques are presented here and so there is no explanation. Remember, in the R environment each new plot that is generated overwrites the previous plot so if you run the entire script at once you will only see the last plot generated. If R studio is used all plots are not overwritten as they are generated.

### Script with Uncompetitive, Competitive and Mixed inhibition

~~~
# Fit multiple types of inhibition
# input the raw data into a data frame
conc <-
c (3.9,1.9,0.7,0.5,0.4,0.3,3.9,1.9,0.7,0.5,0.4,0.3,3.9,1.9,0.7,0.5,0.4,0.3,3.9,1.9,0.7,0.5,0.4,3.9,1.9,0.7,0.5,0.4)
rate <-
c(0.19,0.19,0.18,0.14,0.13,0.08,0.18,0.17,0.15,0.13,0.11,0.11,0.2,0.15,0.11,0.1,0.09,0.07,0.16,0.14,0.1,0.08,0.07,0.16,0.13,0.08,0.08,0.07)
inhibitor <-
c (0,0,0,0,0,0,100,100,100,100,100,100,300,300,300,300,300,300,500,500,500,500,500,700,700,700,700,700)
ki.df <- data.frame(conc,rate,inhibitor)
ki.df$inv.conc <- 1/ki.df$conc
ki.df$inv.rate <- 1/ki.df$rate
~~~

~~~
# determine concentrations of inhibitor used in experiment
inhibitor.conc <- unique(ki.df$inhibitor)
~~~

~~~
# create colors for each inhibitor concentration
inhib.color <- rainbow(length(inhibitor.conc))
~~~

~~~
# perform the fittings
kic.nls <- nls(rate ~ (Vmax * conc / (Km*(1+inhibitor/Kic) + conc)),
data=ki.df, start=list(Km=0.5, Vmax=.2, Kic=300))
mixed.nls <- nls(rate ~ (Vmax * conc / (Km*(1+inhibitor/Kic) +
conc*(1+inhibitor/Kiu))), data=ki.df, start=list(Km=0.5, Vmax=.2,
Kic=3 00, Kiu=100))
kiu.nls <- nls(rate ~ (Vmax * conc / (Km + conc*(1+inhibitor/Kiu))),
data=ki.df, start=list(Km=0.5, Vmax=.2, Kiu=100))
~~~

~~~
# generate a summary of the fits
summary(kic.nls)
summary(mixed.nls)
summary(kiu.nls)
~~~

~~~
# extract coefficients - competitive inhibition
kic.Km <- unname(coef(kic.nls)[“Km”])
kic.Vmax <- unname(coef(kic.nls)[“Vmax”])
kic.Kic <- unname(coef(kic.nls)[“Kic”])
~~~

~~~
# extract coefficients - mixed inhibition
mixed.Km <- unname(coef(mixed.nls) [“Km”])
mixed.Vmax <- unname(coef(mixed.nls)[“Vmax”])
mixed.Kic <- unname(coef(mixed.nls)[“Kic”])
mixed.Kiu <- unname(coef(mixed.nls)[“Kiu”])
~~~

~~~
# extract coefficients - uncompetitive inhibition
kiu.Km <- unname(coef(kiu.nls)[“Km”])
kiu.Vmax <- unname(coef(kiu.nls)[“Vmax”])
kiu.Kiu <- unname(coef(kiu.nls)[“Kiu”])
~~~

~~~
# use the values directly in the equation to calculate line of best fit
fit.data <- expand.grid(x=(1:40)/10, inhib=inhibitor.conc)
fit.data$inv.x <- 1/fit.data$x
fit.data$kic.y <-
kic.Vmax*fit.data$x/(kic.Km*(1+fit.data$inhib/kic.Kic)+fit.data$x)
fit.data$mixed.y <-
mixed.Vmax*fit.data$x/ (mixed.Km*(1+fit.data$inhib/mixed.Kic)+fit.data$x*
   (1+fit.data$inhib/mixed.Kiu))
fit.data$kiu.y <-
kiu.Vmax*fit.data$x/ (kiu.Km+fit.data$x*(1+fit.data$inhib/kiu.Kiu))
fit.data$inv.kic.y <- 1/fit.data$kic.y
fit.data$inv.mixed.y <- 1/fit.data$mixed.y
fit.data$inv.kiu.y <- 1/fit.data$kiu.y
~~~

~~~
############ Plot Data and Best Fit #############
# plot lines of best fit - competitive
# generate a blank plot and then plot the raw data
plot(ki.df$conc,ki.df$rate, pch=“”, main=“Competitive”)
for (i in 1:length(inhibitor.conc)) {
   points(subset(ki.df$conc, ki.df$inhibitor==inhibitor.conc[i]),
subset(ki.df$rate, ki.df$inhibitor==inhibitor.conc[i]), col=inhib.color[i], pch=1)
^}^
for (i in 1:length(inhibitor.conc)) {
   lines(subset(fit.data$x, fit.data$inhib==inhibitor.conc[i]),
subset(fit.data$kic.y, fit.data$inhib==inhibitor.conc[i]),
col=inhib.color[i])
^}^
# plot lines of best fit - mixed
# generate a blank plot and then plot the raw data
plot (ki.df$conc,ki.df$rate, pch=“”, main=“Mixed”)
for (i in 1:length(inhibitor.conc)) {
   points(subset (ki.df$conc, ki.df$inhibitor==inhibitor.conc[i]),
subset(ki.df$rate, ki.df$inhibitor==inhibitor.conc[i]),
col=inhib.color[i], pch=1)
^}^
for (i in 1:length(inhibitor.conc)) {
   lines(subset(fit.data$x, fit.data$inhib==inhibitor.conc[i]),
subset(fit.data$mixed.y, fit.data$inhib==inhibitor.conc[i]),
col=inhib.color[i])
^}^
# plot lines of best fit - uncompetitive
# generate a blank plot and then plot the raw data
plot (ki.df$conc,ki.df$rate, pch=“”, main=“Uncompetitive”)
for (i in 1:length(inhibitor.conc)) {
   points(subset(ki.df$conc, ki.df$inhibitor==inhibitor.conc[i]),
subset(ki.df$rate, ki.df$inhibitor==inhibitor.conc[i]),
col=inhib.color[i], pch=1)
}
for (i in 1:length(inhibitor.conc)) {
   lines(subset(fit.data$x, fit.data$inhib==inhibitor.conc[i]),
subset(fit.data$kiu.y, fit.data$inhib==inhibitor.conc[i]),
col=inhib.color[i])
^}^
~~~

~~~
############ Plot Residuals #############
# look at residuals and plot
ki.df$kic.resid<- resid(kic.nls)
ki.df$mixed.resid<- resid(mixed.nls)
ki.df$kiu.resid<- resid(kiu.nls)
# plot Residuals - competitive
# generate a blank plot and then plot the raw data
plot (ki.df$conc,ki.df$kic.resid, pch=“”, main=“Residuals - Competitive”)
for (i in 1:length(inhibitor.conc)) {
   points (subset (ki.df$conc, ki.df$inhibitor==inhibitor.conc[i]),
subset(ki.df$kic.resid, ki.df$inhibitor==inhibitor.conc[i]),
col=inhib.color[i], pch=1)
^}^
# plot Residuals - mixed
# generate a blank plot and then plot the raw data
plot(ki.df$conc,ki.df$mixed.resid, pch=“”, main=“Residuals - Mixed”)
for (i in 1:length(inhibitor.conc)) {
   points(subset(ki.df$conc, ki.df$inhibitor==inhibitor.conc[i]),
subset(ki.df$mixed.resid, ki.df$inhibitor==inhibitor.conc[i]),
col=inhib.color[i], pch=1)
^}^
# plot Residuals - uncompetitive
# generate a blank plot and then plot the raw data
plot (ki.df$conc,ki.df$kiu.resid, pch=“”, main=“Residuals -
Uncompetitive”)
for (i in 1:length(inhibitor.conc)) {
   points(subset(ki.df$conc, ki.df$inhibitor==inhibitor.conc[i]),
subset(ki.df$kiu.resid, ki.df$inhibitor==inhibitor.conc[i]),
col=inhib.color[i], pch=1)
^}^
~~~

~~~
############ Plot Lineweaver Burk #############
# plot lines of best fit - competitive
# generate a blank plot and then plot the raw data
plot (ki.df$inv.conc,ki.df$inv.rate, pch=“”, main=“Lineweaver Burk –
Competitive”)
for (i in 1:length(inhibitor.conc)) {
   points(subset(ki.df$inv.conc, ki.df$inhibitor==inhibitor.conc[i]),
subset(ki.df$inv.rate, ki.df$inhibitor==inhibitor.conc[i]),
col=inhib.color[i], pch=1)
^}^
for (i in 1:length(inhibitor.conc)) {
   lines (subset(fit.data$inv.x, fit.data$inhib==inhibitor.conc[i]),
subset(fit.data$inv.kic.y, fit.data$inhib==inhibitor.conc[i]),
col=inhib.color[i])
}
# plot lines of best fit - mixed
# generate a blank plot and then plot the raw data
plot (ki.df$inv.conc,ki.df$inv.rate, pch=“”, main=“Lineweaver Burk -
Mixed”)
for (i in 1:length(inhibitor.conc)) {
   points(subset(ki.df$inv.conc, ki.df$inhibitor==inhibitor.conc[i]),
subset(ki.df$inv.rate, ki.df$inhibitor==inhibitor.conc[i]),
col=inhib.color[i], pch=1)
^}^
for (i in 1:length(inhibitor.conc)) {
   lines (subset (fit.data$inv.x, fit.data$inhib==inhibitor.conc[i]),
subset (fit.data$inv.mixed.y, fit.data$inhib==inhibitor.conc[i]),
col=inhib.color[i])
^}^
# plot lines of best fit - uncompetitive
# generate a blank plot and then plot the raw data
plot(ki.df$inv.conc,ki.df$inv.rate, pch=“”, main=“Lineweaver Burk -
Uncompetitive”)
for (i in 1:length(inhibitor.conc)) {
   points (subset (ki.df$inv.conc, ki.df$inhibitor==inhibitor.conc[i]),
subset (ki.df$inv.rate, ki.df$inhibitor==inhibitor.conc[i]),
col=inhib.color[i], pch=1)
^}^
for (i in 1:length(inhibitor.conc)) {
   lines (subset(fit.data$inv.x, fit.data$inhib==inhibitor.conc[i]),
subset (fit.data$inv.kiu.y, fit.data$inhib==inhibitor.conc[i]),
col=inhib.color[i])
^}^
~~~

